# Drug repurposing: halogenated salicylanilides inhibit USP8 catalytic activity and ACTH release by pituitary cells

**DOI:** 10.1101/2025.02.26.640120

**Authors:** Thierry Waltrich-Augusto, Magda Mortier, Caroline Barette, Emmanuelle Soleilhac, Laurence Aubry, Marie-Odile Fauvarque

## Abstract

Ubiquitin-specific protease 8 (USP8) has emerged as a crucial regulator of various cellular processes including membrane dynamics and receptor trafficking. Additionally, USP8 gain-of-function variants are frequently observed in Cushing’s disease, a severe condition characterized by a pituitary adenoma and dysregulated adrenocorticotropic hormone (ACTH) secretion. Moreover, USP8 overexpression contributes to resistance to chemotherapy in cancer. Here, we report the identification of Closantel, a halogenated salicylanilide, as a potent and reversible inhibitor of USP8 catalytic activity. We defined a screening approach in buffering condition counteracting the selection of oxidizing compounds and identified Closantel as a potent inhibitor of USP8 among a library of 2,240 FDA-approved drugs and compounds under clinical development. We demonstrated further that Closantel treatment led to a dose-dependent reduction of ACTH secretion by pituitary cells and resulted in the decrease in *POMC* gene expression, the precursor of ACTH, highlighting its potential therapeutic efficacy in Cushing’s disease. In addition to Closantel, we identified several other halogenated salicylanilides, including Niclosamide that is subjected to several clinical investigations in cancer treatment, as inhibitors of both USP8 activity and ACTH secretion by pituitary cells. Our findings underscore the repurposing potential of halogenated salicylanilides in targeting USP8-mediated cellular dysfunctions in Cushing’s disease and cancer.

## Introduction

The covalent linkage of ubiquitin, a highly conserved polypeptide of 76 amino acids, to the first methionine or internal lysine residues within proteins represents an essential post-translational modification in eukaryotic cells, governing protein degradation by the proteasome, as well as influencing protein activity by regulating their assembly into complexes, their subcellular localization or their conformational changes. The linkage of ubiquitin to itself, or its modification by other post-translational modifications such as phosphorylation and acetylation, generates different types of ubiquitin polymers with high structural diversity (Swatek and Komander, 2016) that are selectively recognized by ubiquitin-binding domain of various kinds of effectors contributing to diverse cellular outputs (Komander and Rape, 2012). Protein ubiquitination involves the sequential action of ubiquitin-activating enzymes (E1), ubiquitin-conjugating enzymes (E2), and ubiquitin ligases (E3). Conversely, deubiquitinating enzymes (DUBs) catalyze the hydrolysis of ubiquitin moieties from protein substrates intricately regulating protein fate and activity (Clague et al., 2019; Mevissen and Komander, 2017; Pohl and Dikic, 2019). In the human genome, the DUB family comprises a hundred members divided into seven subfamilies, with the ubiquitin-specific proteases (USPs) forming the largest one (Mevissen and Komander, 2017; Suresh et al., 2020). Due to their role in major cellular processes and their observed deregulation in cancers or other diseases, DUBs are the subject of extensive research aiming at the development of chemical inhibitors (Chan et al., 2023; Colombo et al., 2010; Han et al., 2020; Schauer et al., 2020).

Specifically, USP8 is essential for the sorting of cell surface receptors governing either their recycling to the plasma membrane or their degradation through the lysosomal pathway, mainly acting at the endosomal membrane in this process (Berlin et al., 2010a; Crespo-Yanez et al., 2018; Dufner and Knobeloch, 2019; Jacomin et al., 2016; MacDonald et al., 2014; Niendorf et al., 2007; Wright et al., 2011). USP8 notably controls the ubiquitinated status of the Epidermal Growth Factor Receptor (EGFR) and of several components of the ESCRT endocytic machinery (Alwan and van Leeuwen, 2007; Berlin et al., 2010b; Crespo-Yanez et al., 2018; Mizuno et al., 2005; Niendorf et al., 2007).

Pioneer genetic sequencing projects and further independent cohort analyses have revealed that gain-of-function variants in USP8 are expressed in 30 to 60 percent (depending on the cohorts) of patients with Cushing’s disease (Ballmann et al., 2018; Ma et al., 2015; Reincke et al., 2015; Simon and Theodoropoulou, 2022; Theodoropoulou et al., 2015), a rare disease characterized by a pituitary tumor (corticotroph adenoma) responsible for the dysregulation of adrenocorticotropic hormone (ACTH) secretion (Lonser et al., 2017). This dysregulation results in a permanent release of cortisol by the adrenal gland cortex, causing various morbidities including central obesity, skin atrophy, muscle wasting, cardiovascular disease, diabetes, metabolic syndrome, osteoporosis, disturbed mood, and impaired reproductive function. These manifestations collectively define the Cushing’s syndrome that may also results from extra-pituitary tumor cells secreting ACTH or an adrenocortical tumor secreting corticoid hormones (Barbot et al., 2020). USP8 mutations identified in the context of Cushing’s disease are almost exclusively clustered in a region upstream of the catalytic domain that binds the 14-3-3 adaptor protein (LKRSYS^718^SP^720^), a negative regulator of USP8 activity. These mutations are dominant possibly inducing a constitutive USP8 activation and deregulation of cellular homeostasis (Bujko et al., 2019; Dufner and Knobeloch, 2019).

Significantly, numerous studies have demonstrated that silencing the USP8 encoding gene or inhibiting USP8 catalytic activity by chemicals leads to a reduction in ACTH secretion by mouse pituitary cells (Fauvarque et al., 2016; Jian et al., 2016; Perez-Rivas et al., 2017; Treppiedi et al., 2021) Furthermore, USP8 knockdown or chemical inhibition has also shown efficacy in suppressing the growth of various cancer cell types and overcoming resistance to chemotherapeutic compounds (Byun et al., 2013; Colombo et al., 2010; Fauvarque et al., 2016; Jeong et al., 2017; Jian et al., 2016; Oh et al., 2014; Rong et al., 2020; Sun et al., 2020; Tian et al., 2024; Treppiedi et al., 2021; Uzilov et al., 2021; Zhu et al., 2021). Hence, USP8 has emerged as a promising therapeutic target for Cushing’s disease patients who are not cured by surgery and harboring *Usp8* gain-of-function mutations or for patients exhibiting overexpressed USP8 and resistance to chemotherapy in cancer.

The catalytic activity of ubiquitin-specific proteases relies on a conserved cysteine residue that is highly susceptible to oxidation by reactive oxygen species (ROS) (Brnjic et al., 2014; Cotto-Rios et al., 2012; Lee et al., 2013; Tyagi et al., 2021) (Ohayon et al., 2015; Snyder and Silva, 2021; Turell et al., 2020). Redox cycling compounds, capable of generating micromolar concentrations of hydrogen peroxide (H2O2) in the presence of strong reducing reagents like dithiothreitol (DTT) can actually inhibit the catalytic activity of the deubiquitinases by oxidizing accessible cysteine residues (Johnston, 2011). Notably, the oxidation of USP2, a ubiquitin-specific protease closely related to USP8, occurs in the presence of quinone-containing compounds and DTT (Gopinath et al., 2017; Gopinath et al., 2016)(M. Mortier, M.-O. Fauvarque and S. Brugière, unpublished observation).

In this study, we performed a screen to identify new USP8 inhibitors in a library of drugs (2140 molecules) approved by the FDA or in clinical trials in a drug repurposing strategy. To avoid the selection of compounds likely to act by oxidation of the catalytic cysteine, we developed buffer conditions in which DTT was replaced by L-Cysteine to significantly diminish redox cycling without preventing DUBs catalytic activity. In these conditions, we identified Closantel, a halogenated salicylanilide, as a robust inhibitor of USP8 catalytic activity *in vitro* and of both ACTH secretion and *POMC* gene expression by pituitary cells. Our results highlight the potential of repurposing Closantel, Niclosamide and other halogenated salicylanilides as USP8 inhibitors as a therapeutic avenue for Cushing’s disease and cancer treatment.

## Materials and Methods

### Chemicals

PCR6236 was kindly provided by our collaborators Pr. P. Vanelle and Dr. V. Remusat from Aix-Marseille University (Fauvarque et al., 2016). The inhibitor DUB-IN-2 (HY-50737A), the Closantel (HY-17596), the Niclosamide (HY-B0497), the Rafoxanide (HY-17598) the Oxyclozanide (HY-17594) and the salicylanilide bacbone (HY-B1408) were purchased from MedChemExpress and solubilized in pure DMSO (10mM stock solution). Other compounds were purchased from Ambinter or MolPort supplier as indicated in Suppl. Table 1. Search for analog was done using Mcule (https://mcule.com/) “1-CLICK SCAFFOLD HOP” tool, with Closantel as input molecule and the Mcule “10k diverse subset of Purchasable compounds” proprietary library as the chemical space. The top 4 molecules that were available “in-stock” from suppliers were acquired to assay USP8 inhibition and ACTH secretion by AtT-20 cells.

### Monitoring USP8^CD^ activity in L-Cysteine versus DTT containing buffer

The in-house catalytic domain of USP8 (USP8^CD^) was purified using standard procedures and was used at a concentration of 80 nM in the presence of the artificial substrate Ubiquitin-Rhodamine 110 (Ub-Rho) (ref: 60-0117-050 Ubiquigent) at a final concentration of 0.1 µM, in conditions ensuring linearity of enzyme activity throughout the assay incubation time. Enzymatic assay was performed in a Tris-HCl buffer (40 mM Tris-HCl pH 7.4, ovalbumin 0.5mg/ml, L-Cysteine-HCl to a final con 5.5 mM). Compounds were added to the reaction mix and incubated with enzyme for 20 min, before addition of the substrate. The enzymatic reaction kinetics were monitored by measuring fluorescence elicited by the hydrolysis of the peptidyl Ub-Rho over 30 min. The negative control was the solvent DMSO at 1% (maximal concentration used in this study).

### Automated Screening for inhibitors of USP8^CD^ catalytic activity

The screening procedure was based on an automated enzymatic assay using ubiquitin molecule conjugated to fluorescent leaving group substrate (i.e., ubiquitin-7-amido-4-methylcoumarin hereafter designated as Ub-AMC, Ubiquigent ref. 60-0116-050) used at a final concentration of 1 µM, in a 96-well format. The in-house purified catalytic domain of USP8 (USP8^CD^) was used at a concentration of 40 nM. We determined the low values of substrate for which the initial velocity of the enzyme rises linearly with the increasing substrate concentration. The bio-inactive control, showing no inhibition of USP8^CD^, was the solvent DMSO at 0.5 % (corresponds to 0% inhibition). The bioactive control, mimicking the desired inhibitory activity, was the reaction mix without USP8^CD^ (mimicking 100% inhibition). The robustness and reliability of the assay was verified through the calculation of the statistical Z’ factor on a control plate including 40 bioinactive and 40 bioactive controls that reached Z’=0.75 (where 0.5<Z’<1 indicates robust assay (Zhang et al., 1999)). Primary screening was performed in simplicate at a final compound concentration of 50 µM in DMSO 0.5%. Compounds were added to the reaction mix just before the substrate. The USP8^CD^ catalytic activity was monitored both before adding the substrate (T0) and after two hours of incubation at room temperature (Final Time TF).

Data analysis was done using the in-house TAMIS software (http://www.bge-lab.fr/Pages/CMBA/TAMIS.aspx): raw data from each compound-containing well were normalized using the mean values of bioactive and bio-inactive controls (100% and 0% inhibition, respectively) to obtain a percentage of inhibition activity for each compound value. The percentage of inhibition was calculated as follows: (sample raw value - bio-inactive controls mean value)*100)/(bioactive controls mean value - bio-inactive controls mean value).

### Hit validation

For hit validation, USP8^CD^ (in-house purified) or full length USP8 (BostonBiochem ref.E-520) were used at a concentration of 40 nM or 75 nM, respectively, in the presence of another ubiquitin substrate, i.e., the Ub-Rhodamine 110 (Ub-Rho) (Ubiquigent ref. 60-0117-050) at a final concentration of 0.1 µM. Since fluorescence emitted by Rhodamine (excitation peak at 546 nm and an emission peak at 568 nm) is different than that emitted by AMC (excitation peak at 341 nm and an emission peak at 441 nm), this rules out an artefact due to direct inhibition of fluorescence emission. Compounds were added to the reaction mix just before the substrate as described above. The enzymatic reaction kinetics were monitored by measuring fluorescence elicited by the hydrolysis of the Ub-Rho bound over 30 min using the same controls as for Ub-AMC in the presence of various doses of each compound. Compounds showing at least 30% inhibition of USP8^CD^ at 5 µM were considered as efficient hits.

### Selectivity assays against other cysteine proteases

Selectivity towards USP8 compared to other DUBs was assessed by monitoring the IC_50_ of Closantel towards full length USP8 used at 75 nM (BostonBiochem ref.E-520) *versus t*he closely similar USP2 enzyme used at 16 µM (TEBU-Bio 80352 or Ubiquigent 64-0014-050) following manufacturer’s instructions and a similar procedure as described for USP8^CD^. Each enzyme was incubated in the presence of Ub-Rho (Ubiquigent ref. 60-0117-050) at a final concentration of 0.1 µM in conditions ensuring linearity of enzyme activity throughout the assay in a specific buffer. Compounds were added to the reaction mix just before the substrate. The enzymatic reaction kinetics were monitored by measuring fluorescence elicited by the hydrolysis of Ub-Rho over 30 min at concentration of Closantel ranging from 100 to 0.1 µM each in triplicate.

### Reversibility assay

The reversibility of Closantel inhibitory effect on USP8 was tested using a gel filtration assay as follows: USP8 at 50 nM was incubated for 15 min with Closantel at 25 µM or with DMSO and passed through a gel filtration column (Millipore ref. UFC30GV25). The enzymatic reaction kinetics was then monitored by measuring fluorescence elicited by the hydrolysis of the Ub-Rho bound over 30 minutes.

### Monitoring ACTH release by pituitary cells and cell viability by Prestoblue^TM^

Mouse pituitary tumor AtT-20 (AtT-20/D16v-F2) cells were obtained from the American Type Culture Collection (ATCC® CRL-1795) and maintained in DMEM containing 10% fetal bovine serum, 100 μg/mL streptomycin and 100 U/mL penicillin at 37°C with 5% CO2. An enzyme immunoassay (EIA) kit was adapted to the automated robotic platform in a 96-well format. 1.25 × 10^4^ AtT-20 cells were seeded in poly-L-lysine-coated 96-well culture plates in 150 µL of medium. After 24 hours, cells were treated with fresh medium containing stock compounds solubilized in DMSO at indicated concentrations or with DMSO alone at 0.5 %. After 24h of treatment, cell culture supernatant was collected, cleared for 10 min at 1000 x *g* at 4°C, diluted in assay buffer (Phoenix Peptide ACTH EIA EK-001-21) and frozen at - 80°C for subsequent processing according to the instructions of the kit’s manufacturer.

After collection of the supernatants for the ACTH assay, cells were supplemented with the PrestoBlue™ cell viability reagent (Invitrogen™ A13261) at 10% final volume, and re-incubated at 37°C for 30 minutes. Fluorescence was then read with a BMG Labtech’s FLUOstar OPTIMA microplate reader (excitation/emission 544nm/580 nm) or a Tecan’s Infinite M1000 (excitation/emission 560 nm/590 nm).

### Live cell monitoring of toxicity on salicylanilides on pituitary cells

AtT-20 cells were seeded at 35,000 cells per well in 96-well plates. After 24h, cells were treated with Propidium Iodide (Sigma Aldrich P4864) at 0.5 µg/ml and compounds at the indicated concentrations, i.e., Staurosporine at 1.0 µM, Camptothecin at 5.0 µM, Closantel and its analogous at 6.25 µM, 12.5µM or 25.0 µM). Plates were then installed in the IncuCyte Zoom® microscope (Sartorius) for automated image acquisition of three images per well in phase contrast and in the red channel at 10x magnification every four hours until 68h. Image analysis of the phase and red channel images allows extracting respectively the % of confluence and PI positive (PI^+^) dead cell counts. From the kinetic curves of each feature, the area under the curve (AUC) were calculated with GraphPad Prism 10 software.

### RT-Q-PCR

To analyze POMC gene expression, 3.75 × 10^5^ AtT-20 cells were seeded on polylysine-coated 6-well culture plates in 2.4 mL complete medium. After 24h, cells were treated with indicated compound diluted in DMSO (DMSO at 0.5% final concentration). After 24h of treatment, cells were briefly rinsed with DPBS (no calcium, no magnesium modification) pre-warmed at 37°C, lysed with the LBP reagent from the NucleoSpin® RNA Plus kit (Macherey-Nagel 740984.50) and then frozen at -70°C before continuing the RNA extraction protocol according to the manufacturer’s instructions. One µg of RNA was used to synthesize cDNA using the AffinityScript QPCR cDNA Synthesis Kit (Agilent #600559). The qPCR reaction was performed using the primers against the mouse POMC gene (5’-CCT CCT GCT TCA GAC CTC CAT AGA T-3’ and 5’-GTC TCC AGA GAG GTC GAG TTT). The expression of the HPRT gene was used as a normalizer (primer pair 5’-ATG GAC AGG ACT GAA AGA CTT GCT-3’ and 5’-TTG AGC ACA CAG AGG GCC ACA ATG-3’). Primers were synthesized by Eurofins Genomics. Detection was done using the Brilliant II SYBR® Green QPCR Master Mix (Agilent #600828), and readings were done using a Stratagene Mx3005P equipment.

### Statistical analysis

Data was processed with the Graphpad Prism 10 software. For both ACTH production and viability assays, data normality was determined by QQ plot inspection. For the ACTH production assay, treatment conditions were compared using a Welch’s ANOVA test followed by a Dunnett’s T3 multiple comparisons test. Data for the viability assay was treated via a Kruskal-Wallis test followed by a Dunn’s multiple comparisons test. For the POMC expression assay, the Ct values of the technical triplicates were first validated via one-sample t-tests for significance using Prism. Normalization to the reference gene and the relative expression levels were then calculated with the Stratagene MxPro v4.0.

## Results

### Screening conditions counteracting the selection of oxidizing molecules

Oxidizing compounds such as chalcone and naphtoquinones were selected as quite potent inhibitors of DUBs (Gopinath et al., 2016; Johnston, 2011; Ohayon et al., 2015) (Oh et al)(Fauvarque et al., 2016). However, concerns have been raised in the field about the potential lack of selectivity and cell toxicity associated with such compounds which can irreversibly oxidize the catalytic cysteine and other reactive residues. To address this issue, we designed buffer conditions substituting dithiothreitol (DTT) with L-Cysteine to minimize or prevent the redox cycle. Under these conditions, we observed that the oxidizing compound PCR6236, a potent irreversible inhibitor of USP8 in the presence of DTT (Fauvarque et al., 2016) (Mortier, M. and Fauvarque, M.O., unpublished results), no longer exhibited inhibitory activity on USP8 catalytic domain (USP8^CD^) (Fig. 1A,B). This experimental modification was next used in our screening conditions to avoid the selection of chemicals that inhibit ubiquitin-specific proteases by irreversibly oxidizing the catalytic cysteine residue.

**Figure 1:**
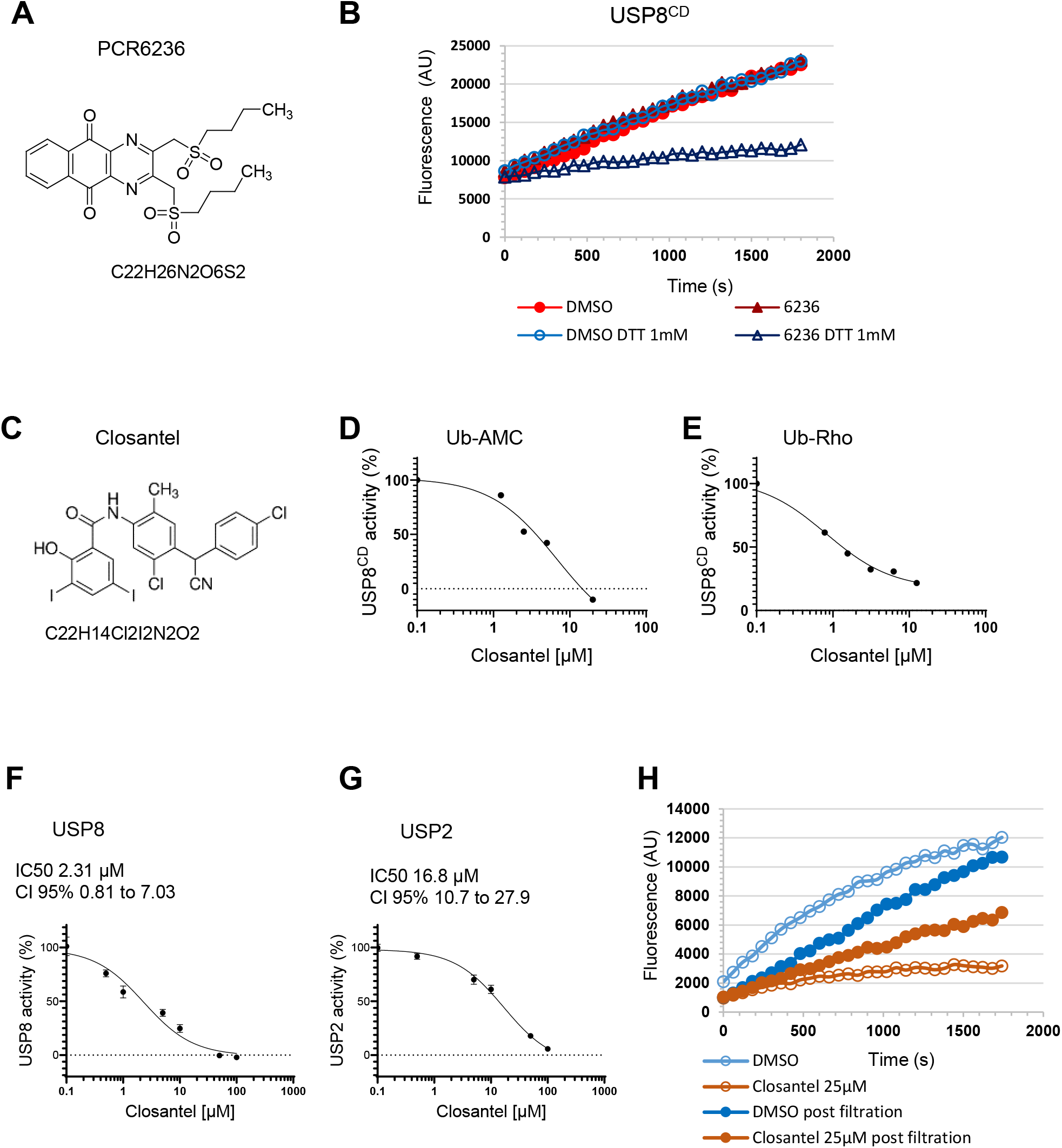
USP8 catalytic activity inhibition by PCR6236 and Closantel. **A**. Chemical structure of PCR6236 compound. **B**. Inhibition of the catalytic activity of USP8 catalytic domain (USP8^CD^) by PCR6236 in the presence of either DTT or L-Cysteine as indicated. DMSO (0.5%) is used as the bio-inactive control resulting in maximal catalytic activity of USP8^CD^. **C**. Chemical structure of Closantel. **D**,**E**. Dose-dependent inhibition of USP8^CD^ by Closantel tested using either Ub-AMC (D) or Ub-Rho (E) as fluorescent substrates, each possessing different emission wavelength. Each experiment is one representative experiment out of two. **F-G**: IC_50_ measurement of USP8 (F), USP2 (G) or UCH-L3 (H) inhibition by Closantel. Error bars indicate standard deviation between triplicates. Calculated IC_50_ and confident interval (CI) are indicated. **H**. USP8 inhibition by Closantel at a 25 µM concentration before and after gel filtration as indicated.

### Repurposing Closantel as a reversible inhibitor of USP8 catalytic activity

We used the Prestwick Chemical Library® (1,280 FDA-approved drugs or compounds under clinical development) and an in-house library (including 960 additional non-redundant FDA-approved drugs selected from Tebu-bio catalog) to identify potential USP8 inhibitors with high biocompatibility. Screening the combined library of 2,240 compounds at a final concentration of 50 µM, we identified seven hits that exhibited robust inhibition of Ub-AMC cleavage at both 50 µM and 5 µM concentrations on USP8^CD^, among which Closantel (Fig. 1C). Dose-dependent analyses confirmed the ability of Closantel to inhibit USP8-dependent cleavage of Ub-AMC as well as of Ub-Rho (Ub-Rhodamine 110), a second substrate with different excitation and emission wavelengths (Fig. 1D,E). Inhibition of USP8-dependent hydrolysis of two different ubiquitin substrates indicates that Closantel is unlikely interfering with fluorescent signals in a non-specific manner.

Additional dose-dependent assays showed that Closantel displayed an IC_50_ of 2.31 µM (95% CI 0.81 to 7.03) against the full-length USP8 (Fig. 1F) and an IC_50_ of 16.8 µM (CI 95% 10.7 to 27.9) against the closely related enzyme USP2 (Fig. 1F-G).

The reversibility of USP8 inhibition by Closantel was assessed using a gel filtration assay. The results indicated a recovery of up to 60% of the control USP8 activity after gel filtration, suggesting that Closantel did not irreversibly inhibit USP8 catalytic activity (Fig. 1H). This finding is consistent with the experimental condition, which were designed to prevent the selection of irreversible redox cycling compounds.

### Halogenated salicylanilides inhibit USP8 catalytic activity

Closantel is a halogenated salicylanilide that shares a salicylanilide backbone with other compounds in the family (Fig. 2, suppl. Table 1). Currently employed as a veterinary antihelminthic drug, Closantel’s usage aligns with other members of this compound class, as discussed below. Notably, this family includes Niclosamide, a compound subjected to several clinical assays for cancer treatment (Barbosa et al., 2019). We next analyzed the effect of Closantel and Niclosamide as well as of a comprehensive set of other halogenated salicylanilides on the catalytic activity of full-length USP8. In addition to Closantel, all ten halogenated salicylanilides tested displayed inhibition of USP8 catalytic activity (Fig. 2, Suppl. Table 1). In contrast, the salicylanilide backbone was mostly inactive when used at concentrations below 100 µM showing only a slight inhibitory effect at 50 µM (Fig. 2, Suppl. Table 1). Employing the F-trees similarities search program on the Mcule platform (https://doc.mcule.com/doku.php?id=ftrees), we selected six additional analogs to Closantel. The activity of these six analogs was assessed on USP8, revealing that none of these compounds inhibited USP8 catalytic activity when tested at 40 µM (Suppl. Table 1). Overall, these data indicate that the active compounds are of the general formula shown in Figure 3. Collectively, our findings show that Closantel and other halogenated salicylanilides effectively inhibit USP8 catalytic activity.

**Figure 2:**
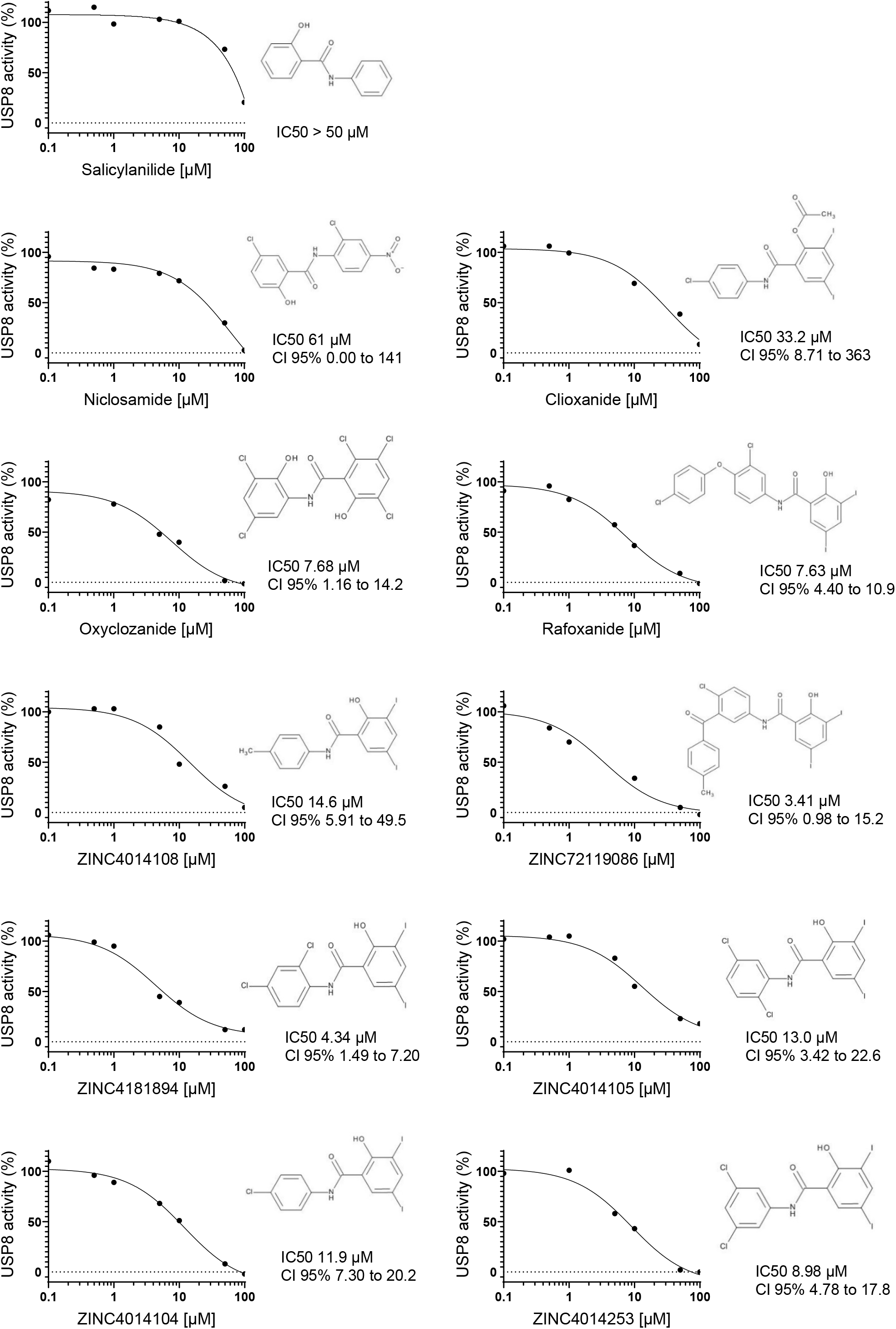
Dose-dependent inhibition of USP8 by salicylanilides. Inhibition of USP8 catalytic activity was monitored using Ub-Rho substrate in the presence of each compound, as indicated, tested at six to seven doses (from 0.1 µM to 100 µM) plus DMSO 0.1% condition, each in simplicate. Calculated IC_50_ and confident interval (CI 95%) are indicated.

**Figure 3:**
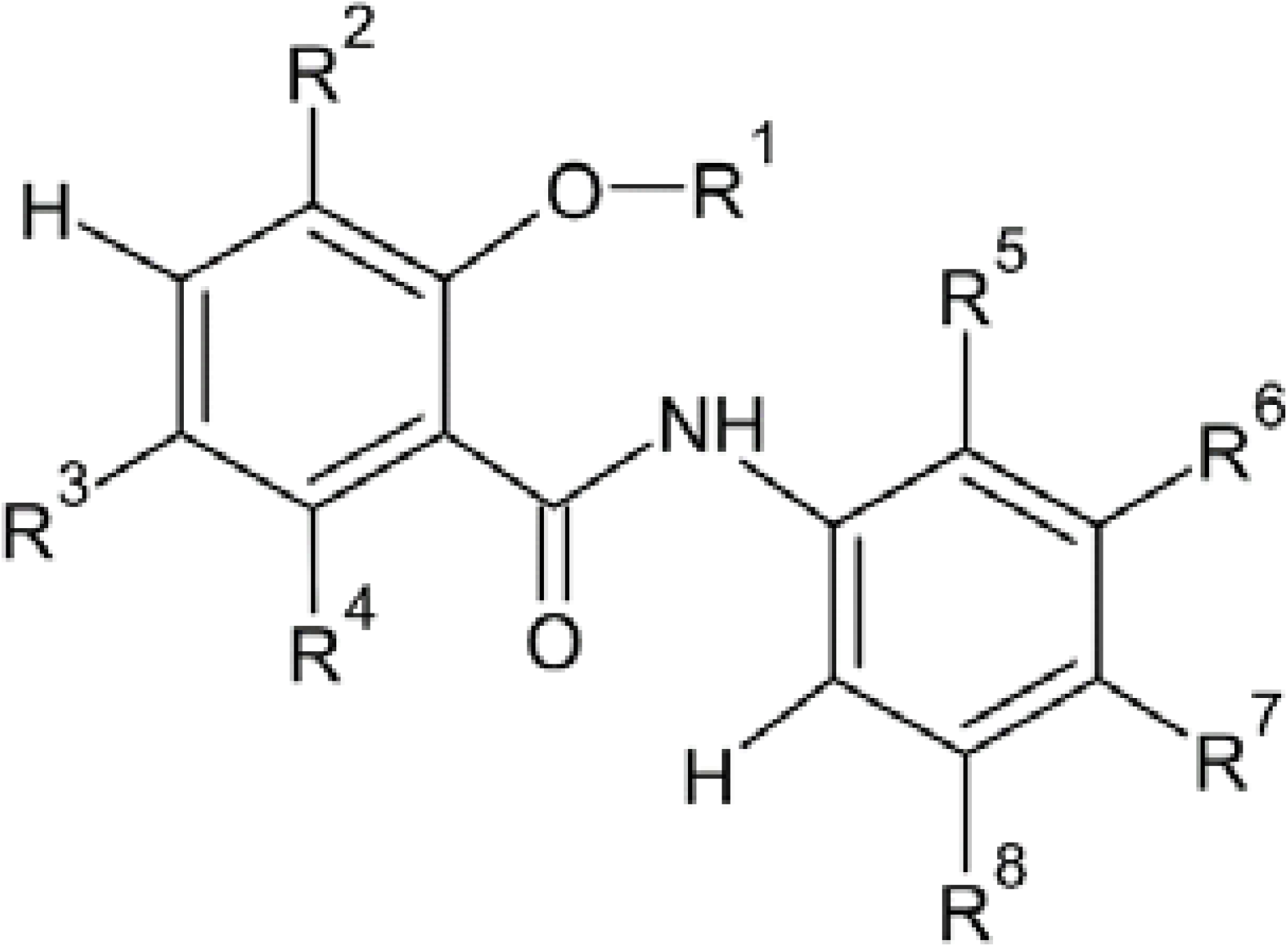
General formula of active compounds inhibiting USP8 catalytic activity. General formula wherein R^1^ is H or a -C(O)-R^9^, wherein R^9^ is a C_1_-C_3_ linear or branched, saturated or unsaturated, alkyl group; R^2^ is H or a halogen; R^3^ is a halogen; R^4^ and R^8^ are, independently, H or a halogen, R^5^ is H, a C_1_-C_3_, linear or branched, saturated or unsaturated, alkyl group, OH, or a halogen; R^6^ is H, a halogen, or - C(O)-Ar, wherein Ar is a C_6_-C_10_ aryl group, Ar being optionally substituted with a C_1_-C_3_ linear or branched, saturated or unsaturated, alkyl group; R^7^ is H, a halogen, a C_1_-C_3_, linear or branched, saturated or unsaturated, alkyl group, -NO_2_, or -B-Ar, wherein B is O or -C(CN)-, and Ar is a C_6_-C_10_ aryl group, Ar being optionally substituted with a halogen, wherein if R^2^ is H, then at least one of R^5^, R^6^, R^7^ and R^8^ is not H, or one of its pharmaceutically acceptable salts.

### Halogenated salicylanilides reduce ACTH release by pituitary cells

One therapeutic objective in selecting USP8 inhibitors is to mitigate ACTH production and secretion by pituitary cells. To address this, we employed the mouse AtT-20 pituitary cell line to monitor ACTH concentration in the cell culture media. Our experiments confirmed that a 24-hour exposure of AtT-20 cells to Closantel significantly decreased ACTH release in a dose-dependent manner (Fig. 4A). Simultaneously, we assessed cellular viability of the same batches of Closantel-treated cells to determine non-toxic treatment doses by monitoring cell reducing capacity with PrestoBlue^TM^ (Fig. 4B). We observed that Closantel caused a highly significant 60% reduction in ACTH release at 30 µM, representing the highest non-toxic concentration in these experimental conditions (Fig. 4A,B). On its side, Niclosamide treatment demonstrated potent inhibition of ACTH release at low doses (i.e., 0.5 µM and 1.0 µM), albeit with a significant decrease in cellular viability at doses starting from 3.0 µM (Fig. 4A,B). Collectively, our results highlight Closantel’s and Niclosamide’s efficacy in inhibiting ACTH release in AtT-20 pituitary cells, with no observed cytotoxic effect of Closantel till a treatment concentration of 30 µM included, whereas Niclosamide’s impact on pituitary cell viability required the use of doses above 3.0 µM.

**Figure 4:**
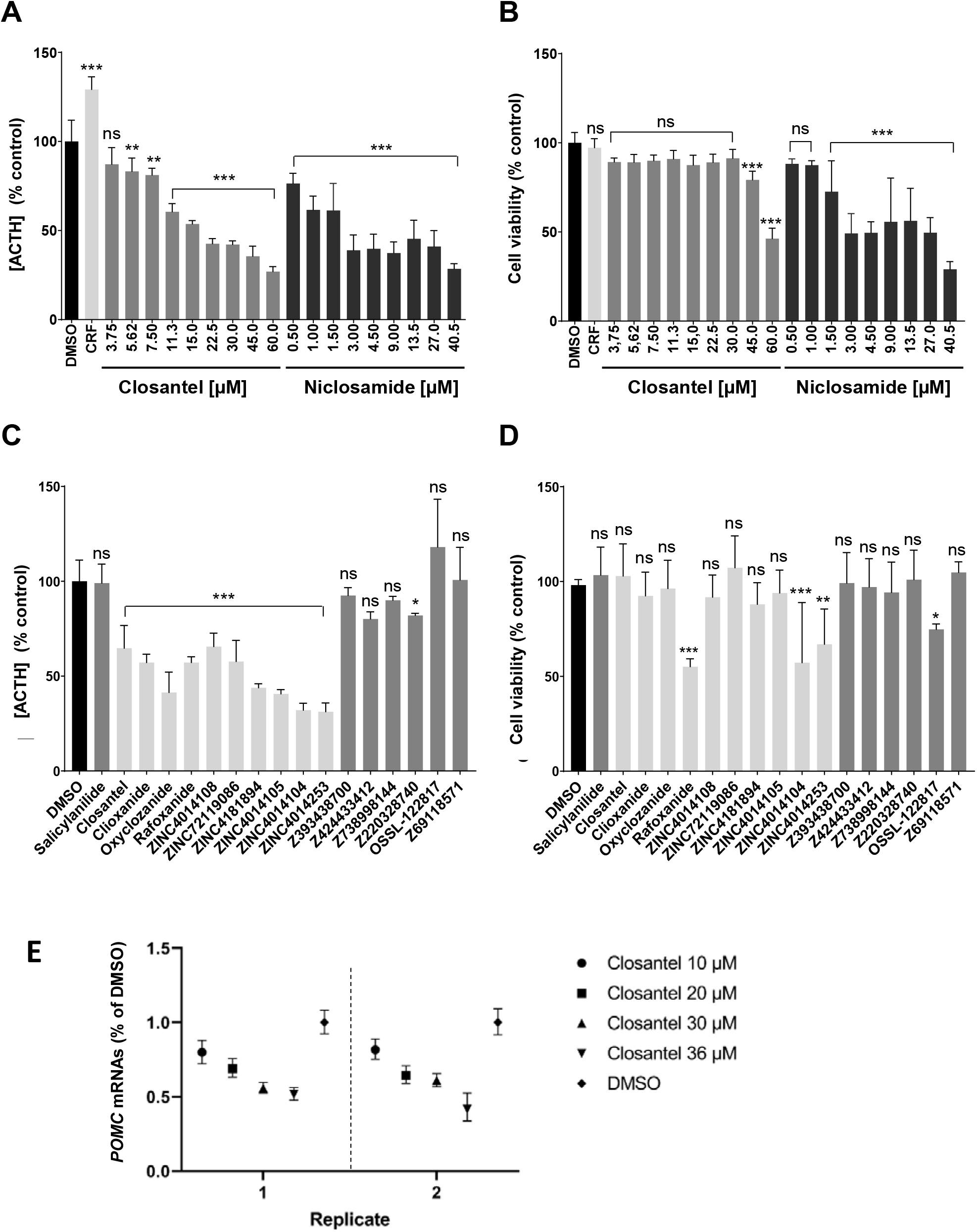
Bioactivity of salicylanilides on AtT-20 pituitary cells. **A,C**. ACTH release in the supernatant of AtT-20 pituitary cells after 24 hours exposure to the solvent DMSO (0.5%), to the ACTH inducer corticotropin-releasing factor (CRF) at 100 nM, to Closantel or Niclosamide at the indicated final concentrations (A) or to each other indicated compound tested at a final concentration of 25 µM (C). Cell supernatants were collected and stored at -80°C, prior to quantification of ACTH concentration using ACTH chemiluminescent EIA kit (Phoenix Pharmaceuticals, Inc.). Determination of ACTH concentrations was based on the best-fitting ACTH standard curve and P values were calculated using one-way ANOVA, using Prism 9 software (GraphPad Software, Inc.). **B, D**. Viability assay of the same AtT-20 cells was concomitantly assessed by using the PrestoBlue™ Cell Viability Reagent that is quickly reduced to a fluorescent product by metabolically active cells. **A-D**. Histograms represent mean values expressed as a percentage of the DMSO control. All compounds were tested in technical replicates (4 to 8 replicates). Error bars represent standard deviation to the mean. **P<0.002. ***P< 0.001, ns: non-significant. **E**. RT-Q-PCR analysis of *POMC* transcript levels in AtT-20 cell lysates following treatment with indicated doses of Closantel for 24 hours. Each dot represents the POMC transcripts levels normalized to the housekeeping gene HPRT and then compared to the DMSO control condition. Error bars indicate the standard deviation between technical triplicates. Two independent experiments are shown.

Remarkably, each of the halogenated salicylanilides that previously demonstrated activity against USP8 catalytic activity also inhibited ACTH release by AtT-20 cells. Indeed, a 24-hour exposure at 25 µM resulted in ACTH release levels ranging from 66% of the DMSO control (100%) for the less active compound (ZINC4014108) to 31% of the DMSO control for the most active compound (ZINC4014253)(Fig. 4C, light grey histograms, Suppl. Table 1). In contrast, compounds inactive against USP8 catalytic activity did not significantly inhibit ACTH release by AtT-20 cells at 25 µM except in one case (Z220328740), albeit to a level below the 30% inhibition threshold set for this study, categorizing it as inactive (Fig. 4C, dark grey histograms, Suppl. Table 1). Simultaneous monitoring of cell reducing capacity with PrestoBlue^TM^ revealed that, under these treatment conditions, two active compounds, Rafoxanide and ZINC4014104, displayed significant toxicity when applied at 25 µM concentration for 24h (Fig. 4D, light grey histograms). This analysis also indicated that the six additional inactive compounds presented moderate (OSSL-122817) or no reduced mitochondrial metabolism under the assay conditions (Fig. 4D, dark grey histograms).

In summary, our findings demonstrate that Closantel and a set of additional halogenated salicylanilides inhibit both USP8 catalytic activity and ACTH release by pituitary cells at non-cytotoxic doses.

### Closantel inhibits *POMC* gene expression

To determine if the observed reduction in ACTH release could result from changes in the expression of the *POMC* precursor gene, we performed quantitative RT-PCR to assess *POMC* expression in AtT-20 cells treated with varying doses of Closantel. Our results indicate a significant and dose-dependent decrease in *POMC* mRNA levels in response to Closantel treatment, suggesting a negative regulatory impact of Closantel on the upstream signaling pathways controlling the expression of the *POMC* gene (Fig. 4E).

### Closantel and active halogenated salicylanilides analogs effects on AtT-20 cell growth

Having observed some toxicity of two members of the halogenated salicylanilide family (Rafoxanide and ZINC1440104) when applied at 25 µM for 24 hours, we further explored the effect of the active halogenated salicylanilides utilized in this study. To this end, AtT-20 cells were exposed to each compound at three different concentrations (6.25 µM, 12.5 µM, and 25 µM) and growth and viability of cells were tracked over a period of 68 hours through automated imaging by monitoring cell confluence and the number of propidium iodide-positive (PI^+^) dead cells respectively. This comprehensive analysis unveiled distinct patterns of cell growth depending on treatment (Fig.5, Suppl. Fig. 1). Notably, cells treated with Closantel displayed a growth rate and PI incorporation similar to those observed in DMSO-treated control cells (depicted at 24h, 48h, and 68h in Fig. 5A) except a modest increase of PI incorporation at 25 µM when assessed over the 68 hours period (Fig. 5A,B). Compounds other than Closantel exhibited varied effects on cell growth and cell death induction with, in a few cases, a lack of strict correlation between the previously monitored metabolic activities compared to the cell confluence analysis. In this setup, Rafoxanide treatment in particular did not induce neither reduced cell growth nor enhanced PI incorporation while it did provoke reduced metabolic activity when tested with Prestoblue^TM^. This suggests that monitoring cell reducing capacity can reveal defective metabolism preceding or independent of cell growth defects. Of particular relevance to our study, this in-depth analysis revealed a notable lack of cytotoxicity for Closantel in the AtT-20 cell culture conditions used in this analysis.

**Figure 5:**
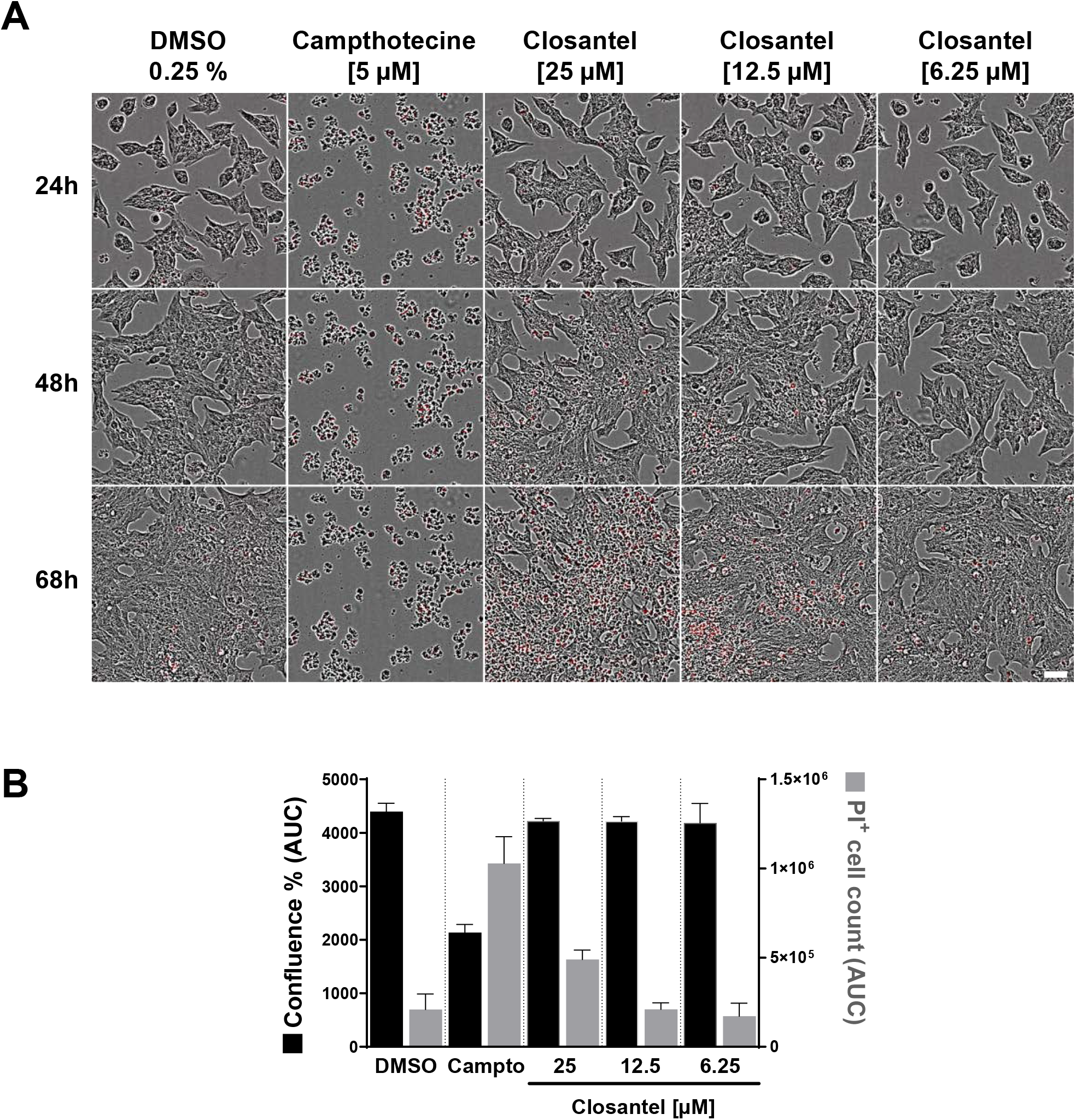
Toxicity evaluation of Closantel on AtT-20 pituitary cells by live cell imaging. **A**. Merge images of phase contrast and PI staining (red) of AtT-20 cells treated with DMSO, Camptothecin (Campto) or Closantel at the indicated concentrations at 24h, 48h and 68h. Image montage was made thanks to the ImageJ plugin FigureJ (Mutterer and Zinck, 2013). Scale bar : 50 µm. **B**. Live cell imaging was conducted during 68h for each compound at 3 concentrations in 3 technical replicates and analyzed to monitor cell growth (confluence) and toxicity (PI^+^ dead cells). Histograms represent the mean of the AUC calculated from the full kinetics over 68h for confluence % (in black, left Y axis) and PI^+^ cell count (in grey, right Y axis) features. Error bars represent standard deviation to the mean.

## Discussion

Cushing’s disease is a rare disease characterized by dysregulated cortisol production due to ACTH-secreting pituitary adenomas that causes significant metabolic disorders and medical challenge. The expression of gain-of-function of USP8 variants has emerged as a main feature of the disease, providing a new avenue for therapeutic intervention through the use of USP8-specific inhibitors (Byun et al., 2013; Colombo et al., 2010; Fauvarque et al., 2016; Tian et al., 2022; Treppiedi et al., 2021). In this study, we described a new class of USP8 inhibitors belonging to the halogenated salicylanilide family and explored their potential role in ACTH secretion by pituitary cells.

Previous studies had demonstrated the potency of naphthoquinones as DUBs inhibitors, but concerns about their lack of selectivity and potential cell toxicity have been raised with such compounds, particularly due to the oxidizing nature of the quinone moiety (Snyder and Silva, 2021). To address this difficulty, buffer conditions substituting dithiothreitol (DTT) with L-Cysteine were designed to minimize the redox cycle. This modification transformed an oxidizing naphthoquinone compound, PCR6236, from a potent USP8 inhibitor into a non-inhibitory chemical, providing a valuable strategy for selectively inhibiting USP8 in a way distinct from irreversible oxidation.

The screening of FDA-approved drug libraries using this buffer condition led to the identification of Closantel as a robust USP8 inhibitor. Unlike quinone-based compounds, Closantel presented a partially reversible mode of action. Expanding the scope to other salicylanilides, we demonstrated potent inhibition of USP8 catalytic activity by ten halogenated salicylanilides with IC_50_ values ranging from 3.41 µM to 61 µM. In contrast, the salicylanilide backbone was inactive against USP8 except at high concentration (i.e. 100 µM).

Beyond USP8 catalytic inhibition, Closantel and other halogenated salicylanilides induced a dose-dependent reduction in ACTH release from cultured pituitary cells. Importantly, Closantel, Niclosamide and other halogenated salicylanilide tested achieved this effect at non-cytotoxic doses, emphasizing their potential therapeutic value in controlling ACTH release without compromising cellular viability. Further investigations into the molecular mechanisms underlying Closantel effects revealed impairment of *POMC* gene expression that encodes ACTH precursor, suggesting that Closantel acts upstream regulatory signals governing POMC transcriptional activation thereby affecting ACTH production. We cannot exclude that Closantel treatment additionally affects ACTH secretion in the extracellular cell culture medium. These results are coherent with the proposed role of USP8 in the stabilization of plasma membrane receptors such as EGFR that activate a MAPK kinase pathway known to control *POMC* gene activation (Childs et al., 1991; Fukuoka et al., 2011).

Closantel has originally been introduced as an anthelmintic drug interfering with the mitochondrial protons gradient and impairing the ATP synthesis of the parasites (Swan, 1999). While used as an anthelmintic in livestock, Closantel has been associated with human toxicity in a limited reported cases attributed to overexposure (E et al., 2023; Kumar et al., 2019; Tabatabaei et al., 2016). Instances of toxicity notably affected the central nervous system, retina, and optic nerve underscoring the importance of cautious consideration in safety thresholds. Nevertheless, the salicylanilides have been the subject of renewed interest and intensive repurposing studies over the past 10 years, notably in in the search for anti-viral drugs (Jeon et al., 2020; Xu et al., 2020) or anti-cancer treatments (Barbosa et al., 2019; Han et al., 2018; Li et al., 2016; Zhu et al., 2016), a field in which USP8 is also recognized as a valuable therapeutic target to counteract both cell proliferation and resistance to chemotherapies.

## Conclusions

Here, we identified Closantel, Niclosamide and other halogenated salicylanilides as new inhibitors of USP8, a known contributor of Cushing’s disease and cancer resistance to chemotherapy, providing a foundation for future therapeutic inhibitors development within this compound class.

## Author’s contribution

TWA performed ACTH and cell viability assays, RT-QPCR, *in silico* search for analogs and significantly contributed to the discussion of the results ; MM setup screening buffer conditions and performed DUBs activity assays, ACTH and cell viability assays, cell confluence and IP incorporation tests; CB setup the automated USP8 activity assay, performed the robotic primary and secondary screens, interpreted the data for hit selection and supervised automated monitoring of ACTH release; ES supervised the cell confluence and IP incorporation tests; LA co-supervised parts of the study and performed cellular assays; MOF designed the strategy, supervised the study, obtained the funding and wrote the manuscript. All authors contributed to the writing and editing of the manuscript.

## Acknowledgments

The authors thank Dr. M. Balakirev, Dr A. Bouron, Dr. A. Journet, Dr. R. Prudent, Dr. T. Rabilloud, Dr. E. Taillebourg for valuable discussion, information sharing, and technical advices.

## Supplemental material

**Supplementary Table 1**: Formula, structure of chemicals described in the study and of their inhibitory effect on USP8 enzymatic activity, ACTH secretion and AtT-20 cell viability

**Supplementary Fig 1**: **Toxicity evaluation of active halogenated salicylanilides other than Closantel on AtT-20 pituitary cells as monitored by live cell imaging**. Live cell imaging was conducted during 68h for each compound at 3 concentrations in 3 technical replicates and analyzed to monitor cell growth (confluence) and toxicity (PI^+^ dead cells). Histograms represent the mean of the AUC calculated from the full kinetics over 68h for confluence % (in black, left Y axis) and PI^+^ cell count (in grey, right Y axis) features. Error bars represent standard deviation to the mean.

