## Supplemental Table 1 for "Drug repurposing: halogenated salicylanilides inhibit USP8 catalytic activity and ACTH release by pituitary cells"

**Suppl. Table 1. Compounds Activity against USP8 FL, ACTH secretion and AtT-20 cell viability**

| **ZINC number**<br>**Name**<br>**Cat.Number/Ref.**<br>Formula<br>MW | **Chemical Structure** | **USP8FL inhibition**<br>$IC_{50}$ [µM]<br>(95% confidence Interval)<br>(Fig. 1F , Fig.2) | **ACTH secretion**<br>at 25µM<br>*at 27 µM<br>% of control<br>(Fig.4) | **AtT-20 cell viability**<br>at 25µM<br>*at 27 µM<br>% of control<br>(Mean (SD))<br>(Fig.4) |
|---|---|---|---|---|
| **ZINC2057**<br>**Salicylanilide**<br>**MCE HY-B1408**<br>C13H11NO2<br>MW: 213 | 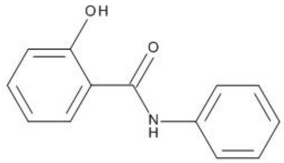 | >50µM | 99% | 105%<br>(14.8) |
| **ZINC4212651**<br>**Closantel**<br>**MCE HY-17596**<br>C22H14Cl2I2N2O2<br>MW: 663 | 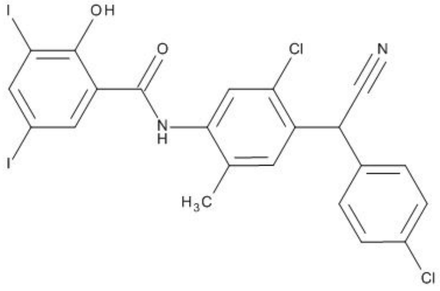 | 2.31 µM<br>(0.81 to 7.03)<br>(Fig. 1F) | 65% | 105%<br>(16.1) |
| **ZINC3874496**<br>**Niclosamide**<br>**MCE HY-B0497**<br>C13H8Cl2N2O4<br>MW: 327 | 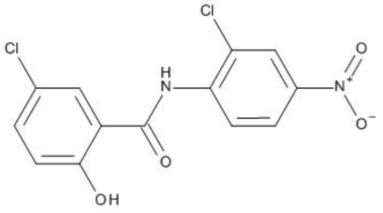 | 61 µM<br>(0.0 to 141)<br>(Fig. 2) | 41%* | 50.0%*<br>(12,4) |
| **ZINC4215387**<br>**Clioxanide**<br>**Amb4466501**<br>C15H10ClI2NO3<br>MW: 541 | 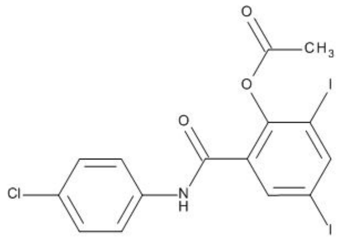 | 33.2 µM<br>(8.71 to 363)<br>(Fig. 2) | 57% | 94.2%<br>(12.3) |
| **ZINC2038733**<br>**Oxyclozanide**<br>**MCE HY-17594**<br>C13H6Cl5NO3<br>MW: 401 | 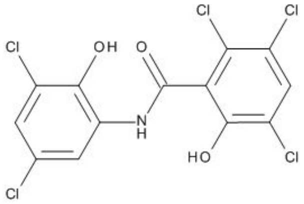 | 7.68 µM<br>(1.16 to 14.2)<br>(Fig. 2) | 41% | 98.1%<br>(14.4) |

| **ZINC number**<br>**Name**<br>**Cat.Number/Ref.**<br>Formula<br>MW | **Chemical Structure** | **USP8FL inhibition**<br>$IC_{50}$ [µM]<br>(95% confidence Interval)<br>(Fig. 1F , Fig.2) | **ACTH secretion**<br>at 25µM<br>*at 27 µM<br>% of control<br>(Fig.4) | **AtT-20 cell viability**<br>at 25µM<br>*at 27 µM<br>% of control<br>(Mean (SD))<br>(Fig.4) |
|---|---|---|---|---|
| **ZINC4181896**<br>**Rafoxanide**<br>**MCE HY-17598**<br>C19H11Cl2I2NO3<br>MW: 626 | 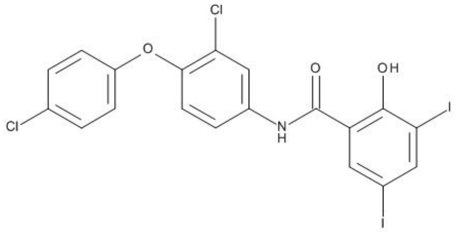 | 7.63 µM<br>(4.40 to 10.9)<br>(Fig. 2) | 57% | 56.2%<br>(3.54) |
| **ZINC4181896**<br>**Rafoxanide**<br>**Amb2704313** | *Idem* | 8.21 µM<br>(3.82 to 18.5)<br>(not shown) | 33% | 39.3%<br>(6.852) |
| **ZINC4014108**<br>**Amb8477380**<br>C14H11I2NO2<br>MW: 479 | 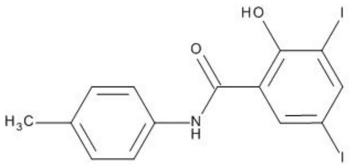 | 14.6 µM<br>(5.91 to 49.5)<br>(Fig. 2) | 66% | 93.5%<br>(11.6) |
| **ZINC72119086**<br>**Amb4103735**<br>C21H14ClI2NO3<br>MW: 617 | 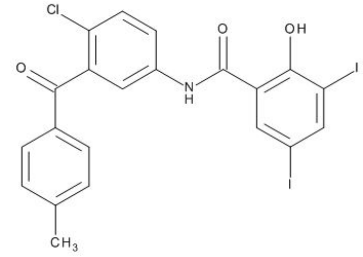 | 3.41 µM<br>(0.98 to 15.2)<br>(Fig. 2) | 58% | 109%<br>(18.2) |
| **ZINC4181894**<br>**Amb8487151**<br>C13H7Cl2I2NO2<br>MW: 533 | 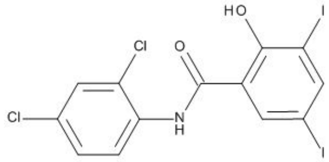 | 4.34 µM<br>(1.49 to 7.20)<br>(Fig. 2) | 44% | 89.6%<br>(10.1) |
| **ZINC4014105**<br>**Amb8477378**<br>C13H7Cl2I2NO2<br>MW 533 | 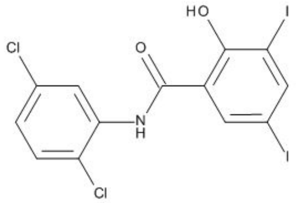 | 13.0 µM<br>(3.42 to 22.6)<br>(Fig. 2) | 41% | 95.7%<br>(11.4) |

| ZINC number<br>Name<br>Cat.Number/Ref.<br>Formula<br>MW | Chemical Structure | USP8FL inhibition<br>$IC_{50}$ [µM]<br>(95% confidence Interval)<br>(Fig. 1F , Fig.2) | ACTH secretion<br>at 25µM<br>*at 27 µM<br>% of control<br>(Fig.4) | AtT-20 cell viability<br>at 25µM<br>*at 27 µM<br>% of control<br>(Mean (SD))<br>(Fig.4) |
|---|---|---|---|---|
| **ZINC4014104**<br>**Amb1935028**<br>C13H8ClI2NO2<br>MW :499 | 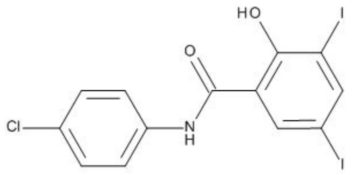 | 11,9 µM<br>(7.30 to 20.2)<br>(Fig. 2) | 32% | 58.8%<br>(34.1) |
| **ZINC4014253**<br>**Amb8477462**<br>C13H7Cl2I2NO2<br>MW : 533 | 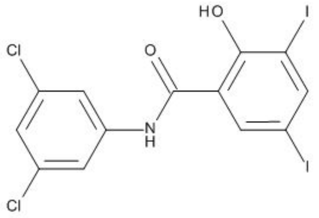 | 8.98 µM<br>(4.78 to 17.8)<br>(Fig. 2) | 31% | 68.0%<br>(17.5) |
| **ZINC32922917**<br>**MolPort**<br>**Z393438700**<br>C20H17FN2O2S<br>MW: 368 | 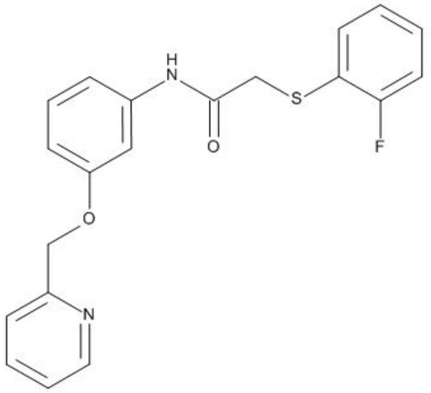 | >40µM<br>(not shown) | 92% | 101%<br>(16.0) |
| **ZINC40111866**<br>**MolPort**<br>**Z424433412**<br>C18H15BrN2O2S<br>MW : 403 | 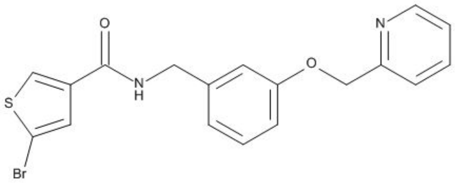 | >40µM<br>(not shown) | 80% | 98.9%<br>(14.7) |

| **ZINC number**<br>**Name**<br>**Cat.Number/Ref.**<br>Formula<br>MW | **Chemical Structure** | **USP8FL inhibition**<br>$IC_{50}$ [µM]<br>(95% confidence Interval)<br>(Fig. 1F , Fig.2) | **ACTH secretion**<br>at 25µM<br>*at 27 µM<br>% of control<br>(Fig.4) | **AtT-20 cell viability**<br>at 25µM<br>*at 27 µM<br>% of control<br>(Mean (SD))<br>(Fig.4) |
|---|---|---|---|---|
| **ZINC55784788**<br>**MolPort**<br>**Z738998144**<br>C21H20N2O3<br>MW : 348 | 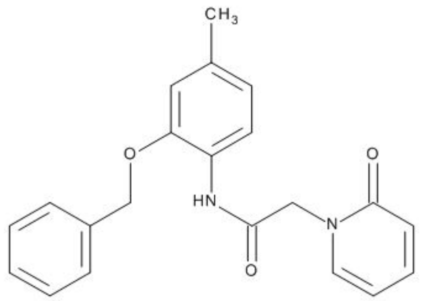 | >40µM<br>(not shown) | 90% | 96.0%<br>(15.33) |
| **ZINC9579366**<br>**MolPort**<br>**Z220328740**<br>C19H12ClFN2O3<br>MW: 370 | 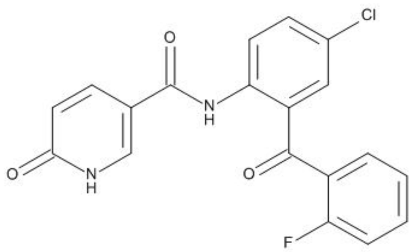 | >40µM<br>(not shown) | 82% | 103%<br>(15.2) |
| **ZINC3159902**<br>**MolPort**<br>**OSSL-122817**<br>C14H12ClNO2<br>MW : 261 | 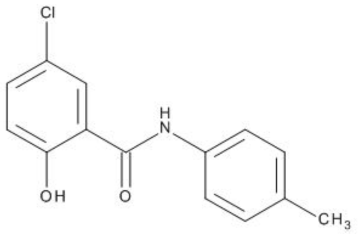 | >100µM<br>(not shown) | 118% | 75.6%<br>(8.67) |
| **ZINC194061705**<br>**MolPort**<br>**Z69118571**<br>C13H10INO3<br>MW: 355 | 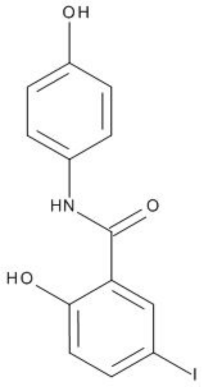 | >100µM<br>(not shown) | 101% | 107%<br>(20.9) |
