## Supplementary figures and images for "Drug repurposing: halogenated salicylanilides inhibit USP8 catalytic activity and ACTH release by pituitary cells"

### Supplemental Figure 1

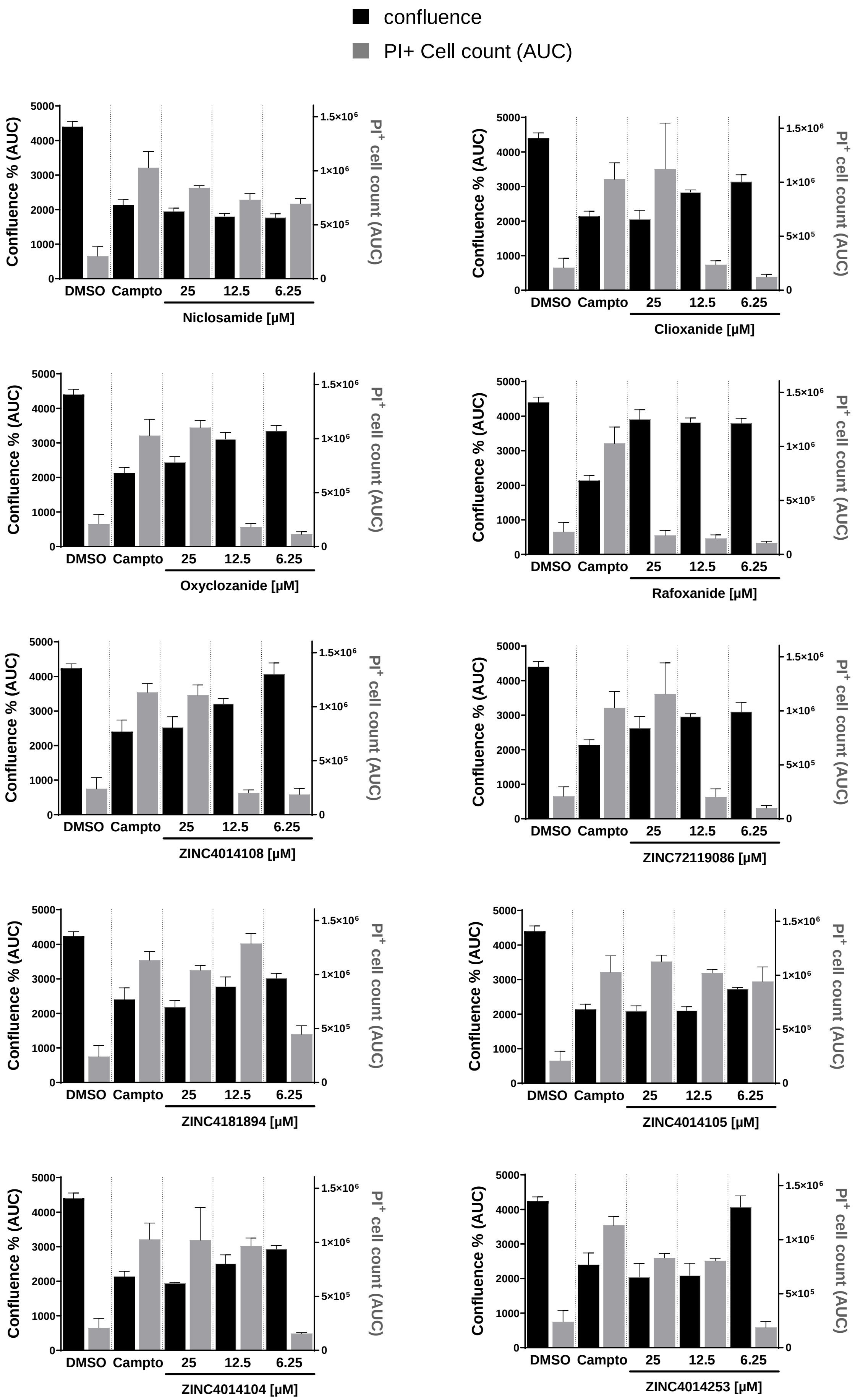


TWA et al., Supplemental Fig 1
